# Effects of Forefoot versus Rearfoot Landing on Biomechanical Risk Factors for Lower Limb Injuries and Performance During Stop-Jump Tasks

**DOI:** 10.1101/2024.12.27.630475

**Authors:** Tianchen Huang, Yuqi He, Lizhi Mao, Mianfang Ruan, Daisuke Takeshita

## Abstract

**Background:** Lower limb injuries commonly occur during sudden deceleration movements. Instructing landing with forefoot or rearfoot is hard to answer, especially considering the injury prevention and keep the performance. Therefore, the purpose of this study is to evaluate the effect of the forefoot and rearfoot landing on the biomechanical risk of lower limb injury prevention and performance during the stop-jumping task.

**Method:** Twenty-three male health subjects were recruited for this study. During a stop-jumping task, three-dimensional kinematic and kinetic, and performance data were collected under two conditions: forefoot landing at initial ground contact and rearfoot landing at initial ground contact. Statistical parametric mapping analysis was used to compare the differences between different landing strategies.

**Result:** Significant differences were found in ankle internal rotation angle and ankle joint moment at foot initial contact with ground between different landing strategies. Landing with forefoot has shorter stance time compared with landing with rearfoot. In the rearfoot strike, posterior ground reaction force (GRF) and GRF inclination angle were smaller than forefoot strike in 0-14% of the stance phase.

**Conclusion:** Landing with forefoot may decrease the risk of non-contact anterior cruciate ligament injuries and have an advancement in quick reaction time, as indicated by decreased stance time, but the risk of lateral ankle sprain may increase for the stop-jump task.

## 1. Introduction

Lower limb injuries are highly prevalent in professional sports, with the knee accounting for 29.3% of such injuries, followed by the ankle (22.4%), thigh (17.2%), and foot (9.1%), collectively posing a significant burden on athletes, individuals, and teams(Mack et al. 2020). Anterior cruciate ligament injuries (ACLIs), especially non-contact ones (NACLIs), account for 70% of all ACLIs (Sanders et al. 2016; Cooke et al. 2003). These injuries not only force athletes to reduce their level of competition but also increase the long-term incidence of osteoarthritis, significantly affecting quality of life, with a substantial risk of re-rupture upon returning to sports (Paterno et al. 2014; Gillquist and Messner 1999).

Ankle sprains, particularly lateral ankle sprains, represent another common category of sports-related injuries (Birrer et al. 1999; Wilkerson and Horn-Kingery 1993). Though often considered minor, ankle sprains can lead to significant muscle and ligament damage, elevate the risk of recurrence, and in severe cases, result in fibular fractures or progress to chronic ankle instability. This can severely impair joint function and pose ongoing health risks (Herzog et al. 2019)(Miklovic et al. 2018).

During physical activities, the moment of landing is considered a critical phase associated with a high risk of lower limb injuries. Proper landing techniques, such as performing a rolling motion when descending from a height, can effectively dissipate impact forces, whereas rigid and vertical landings may lead to catastrophic injuries. Adopting appropriate landing strategies is essential for reducing lower limb joint injuries. Research has identified several biomechanical risk factors related to lower limb injuries are as follows: for ACLIs, including smaller knee flexion angles, greater shear forces, and higher quadriceps forces (Yu and Garrett 2007); for ankle sprains, larger inversion and internal rotation angles, and inversion moments (Kristianslund, Bahr, and Krosshaug 2011). Many studies have evaluated landing techniques based on their potential to mitigate these risk factors (Boyi Dai et al. 2015; Zhou et al. 2021; Chappell et al. 2007). However, recent findings by Boden et al.(Boden and Sheehan 2022) suggest that instead of anterior shear forces causing ACL strain, compressive forces perpendicular to the tibia may play a more significant role in ACLIs.

Considering that injury prevention strategies are often implemented across entire sports teams, evaluating landing techniques concerning risk factors associated with ankle injuries is equally important. Despite this, research on this topic remains limited (Doherty et al. 2014). Additionally, beyond sagittal plane analyses, risk factor evaluations in the frontal and transverse planes should also be incorporated into comprehensive assessments (McLean et al. 2004). To better understand the effects of different landing strategies on ankle and knee injuries, it is crucial to analyze how these strategies influence related ankle and knee joint biomechanical risk factors in three dimensional planes.

Stop-jump maneuvers, which include quick deceleration, landing movement, and jumping vertically, are commonly performed in sports like basketball and volleyball. These maneuvers serve as important technical skills but also carry a high risk of lower limb injuries (Edwards et al. 2012). There are two distinct landing styles: the forefoot strike, which involves making initial contact with the toes, and the rearfoot strike, in which the heel touches the ground first, preceding a stop-jump movement. Research using video analyses of injury incidents reveals that a significant proportion of athletes who sustain ACL injuries during competitions exhibit a rearfoot landing pattern. One study found that different landing techniques influence knee joint kinematics and kinetics during sidestepping maneuvers in female athletes (Cortes et al. 2012). However, movement patterns differ between various landing techniques, and conclusions drawn from sidestepping studies cannot be directly applied to jump-stop maneuvers. Furthermore, most studies on injury risk factors lack an integrated examination of the impact on athletic performance, leaving a critical gap in understanding the balance between injury prevention and performance optimization. This balance becomes particularly critical in stop-jump movements, where large impact forces may increase ankle sprain risk, yet the effects of different landing techniques on both performance and injury risk remain understudied. A better understanding of these mechanisms could lead to more comprehensive training programs that optimize both performance and injury prevention for the knee and ankle.

The purpose of this study was to compare the knee and ankle kinematics and kinetics, and GRF metrics during stop-jump movements using forefoot and rearfoot landing techniques. We hypothesized that compared to rearfoot landing, forefoot landing would result in higher jump height but also increase risk of lateral ankle sprain.

## 2. Method

### 2.1 Participants

The experimental procedure was approved by the University of Tokyo Ethics Committee. Twenty-three healthy, recreational active males were recruited for this study (age: 26.5±8.7 years; height: 1.78±0.05 m; weight:74.84±10.21 kg). Each participant provided written informed consent. To ensure participants standardization, inclusion criteria included the following: (1) no history of lower limb surgical procedures; (2) participants were required to engage in recreational physical activity at least three times per week, with each session lasting a minimum of 30 minutes.

### 2.2 Experimental protocol

Participants were asked to warm up for five minutes in their own manner, which generally consisted of running and stretching. As all participants were right-leg dominant, which were determined based on the preferred jumping leg in a single leg jump for distance, the experiment and data analyses were conducted on the right leg. All participants wore identical models of spandex shorts and shoes (sized appropriately for each participant) as required for the formal experiment. Forty retroreflective markers were placed bilaterally on the acromioclavicular joints, anterior superior iliac spines, posterior superior iliac spines, greater trochanters, medial and lateral femoral epicondyles, medial and lateral malleoli, and first and fifth metatarsophalangeal joints, first toes, and heels (Figure 1). Clusters of four non-collinear markers mounted on rigid thermoplastic shells were attached to the lateral sides of both thighs and shanks using neoprene straps (Cappozzo et al. 1997).

**Figure 1.**
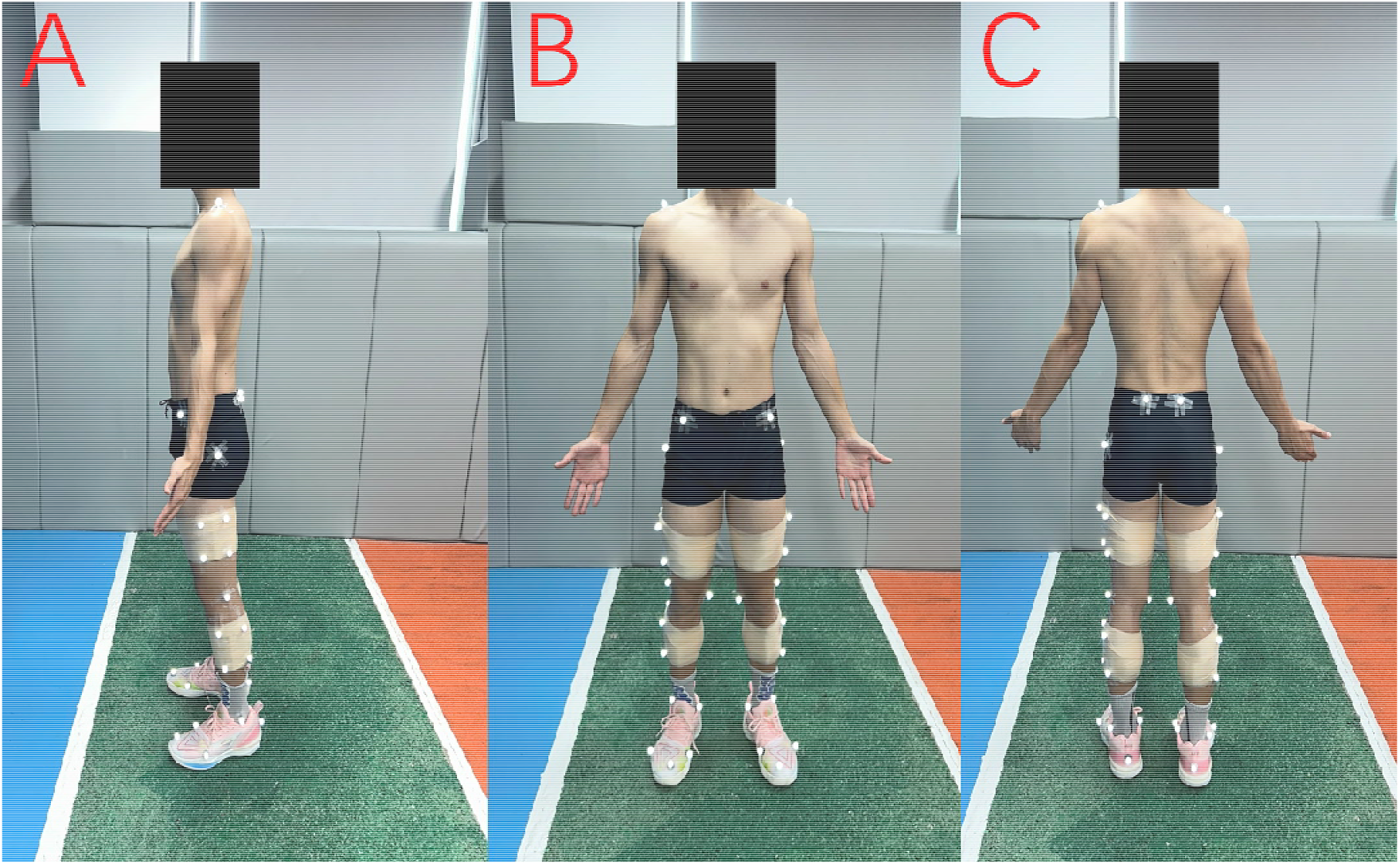
Retroreflective marker placement shown from three views: (A) sagittal, (B) anterior, and (C) posterior. Markers were placed bilaterally on anatomical landmarks, with rigid marker clusters attached to the thighs and shanks using neoprene straps.

A static calibration trial lasting three seconds was conducted, after which the six anatomical markers (medial side of both knees, ankle, and foot) were removed. Once the markers were attached, participants were asked to stand on the force plate to record static data prior to the start of formal data collection. The stop-jump task consisted of an approach run, both-foot landing, and a vertical jump. The participants were instructed to complete the movement as quickly as possible and jump as high as they could. The experimental setup was shown in Figure 2. Five successful stop-jump trials were required in this study. If the participant stepped off the force plate with their right foot or did not perform a vertical jump, they were asked to repeat the trial. As shown in Figure 3, the 2 experimental conditions for the stop-jump task were (1) forefoot landing; (2) rearfoot landing at initial ground contact. The sequence of tasks and the conditions for each task were randomized individually for each participant. This randomization was carried out by sorting random numbers generated using MATLAB. There was 2-minute rest between conditions and a 30-second rest between trials to avoid any fatigue.

**Figure 2.**
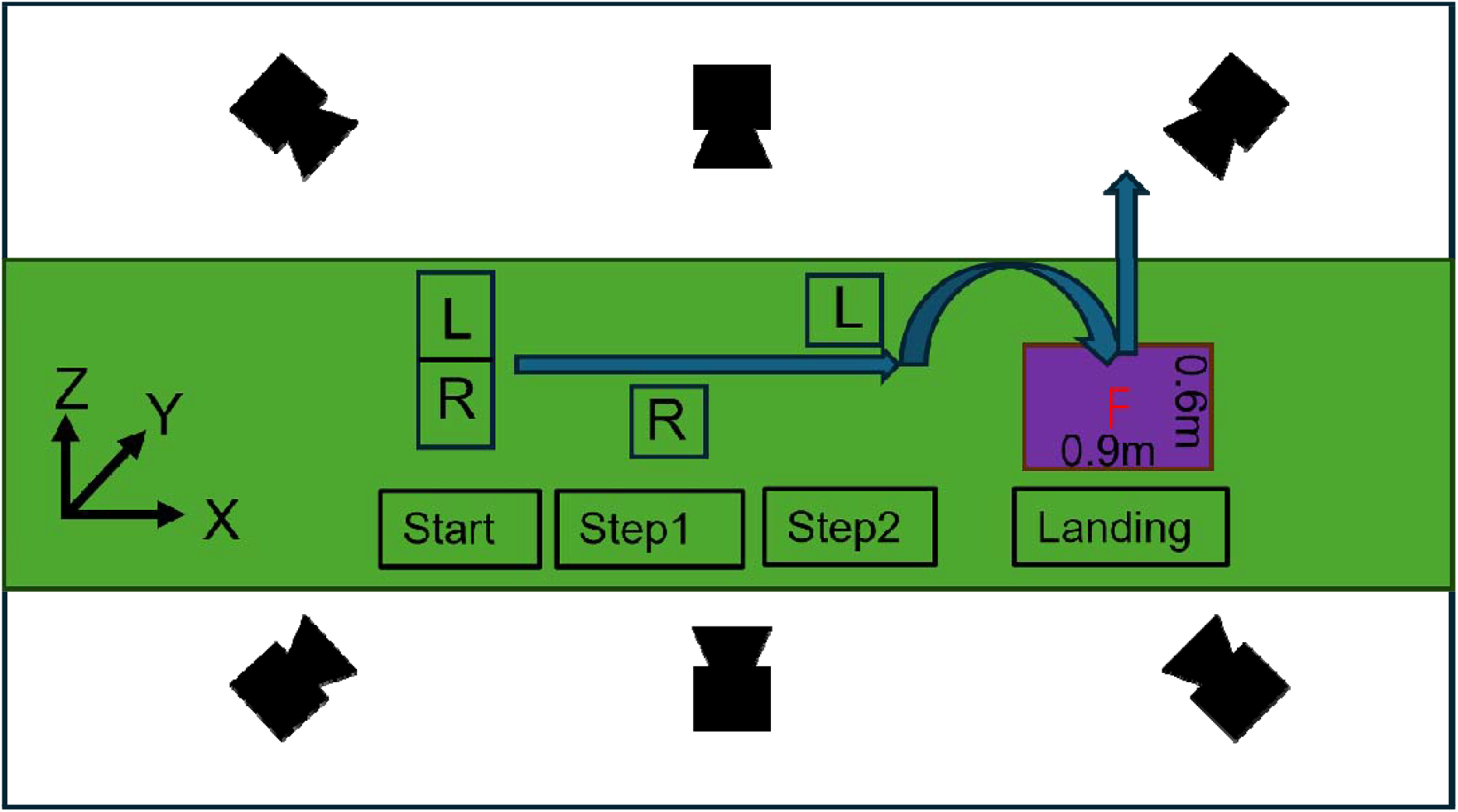
Experimental setup for the stop-jump task. The global coordinate system is defined as: X (anterior+), Y (lateral+), Z (vertical+). The approach path includes start position, and two stepping positions (Step1, Step2) marked for foot placement (L: left foot; R: right foot), leading to a 0.9 m × 0.6 m force platform for the landing phase.

**Figure 3.**
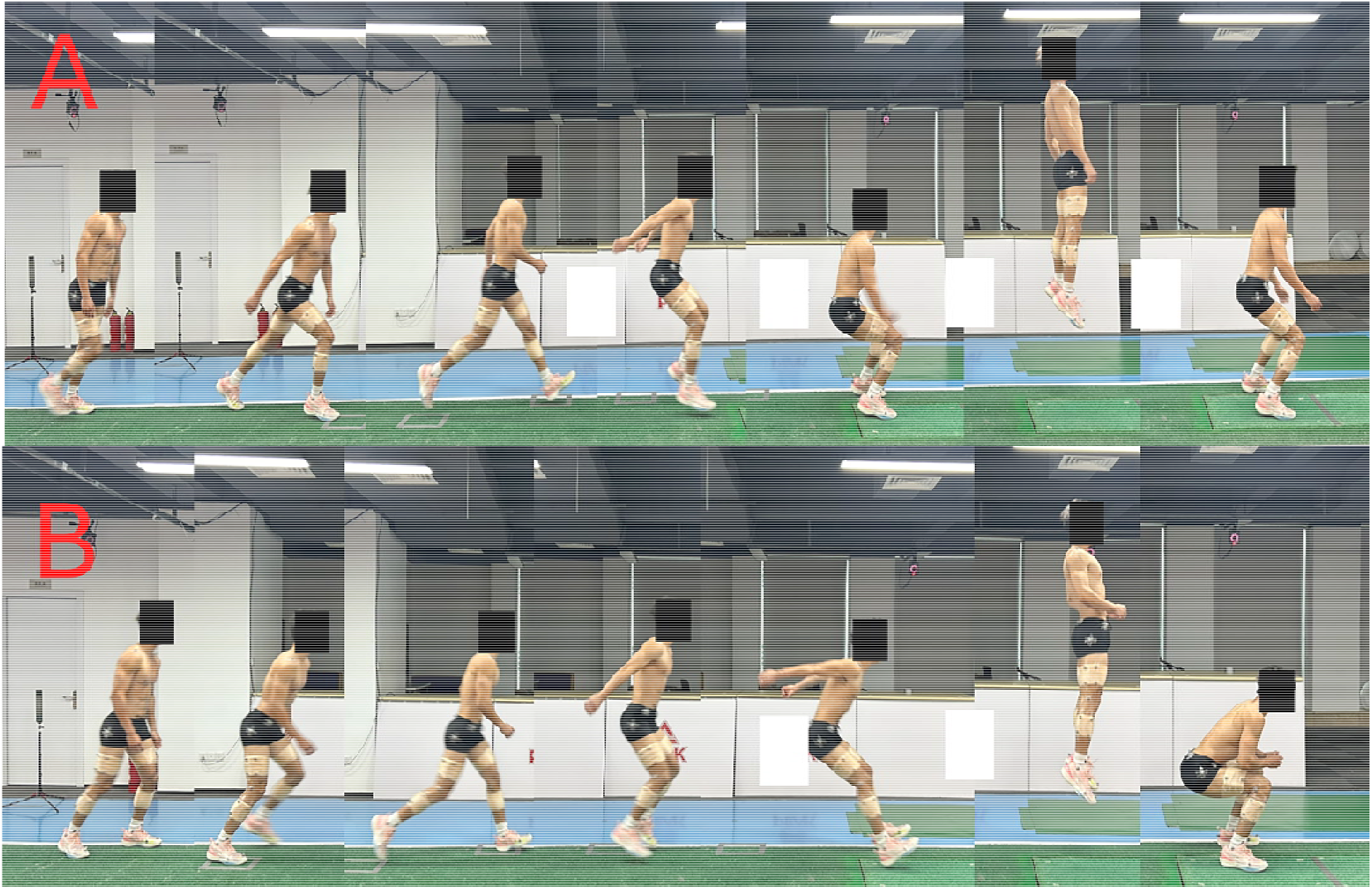
Stop-jump task sequences showing (A) forefoot landing and (B) rearfoot landing techniques. Each sequence illustrates the approach run, landing phase, and vertical jump.

### 2.3 Data collection

A motion capture system consisting of thirteen Mars 4H cameras, 4.1-megapixel resolution (NOKOV Motion Capture System, Beijing, China), was used to collect raw marker coordinate data at 240Hz. A force platform (1200Hz, 9287CA, Kistler Instruments, Winterthur, Switzerland) was placed below ground level in the middle of the runway for ground reaction force data collection. These two data collection systems were synchronized.

### 2.4 Data reduction and processing

The video recordings of the retroreflective marker trajectories were digitized using the NOKOV Seeker video analysis system. For each participant, the five trials of each stop-jump task condition were digitized, starting from 10 frames before right foot initial ground contact to 10 frames after takeoff. To ensure consistency between kinematic and kinetic data under filtering, we applied a low-pass filter with the same cut-off frequency of 6 Hz to both ground reaction force and marker coordinate data. Force-based gait events were used to obtain the time-normalized landing phase. The ground contact phase was defined as the duration from initial contact to takeoff during the first landing of the stop-jump task. Initial contact and takeoff were determined using a vertical ground reaction force (GRF) threshold of 10 N (Zhou et al. 2021; Yu, Lin, and Garrett 2006). The duration of landing phase was scaled to 101 data points. Visual3D biomechanical software (C-Motion, Germantown, MD, USA) was used to create a kinematic model made of seven skeletal segments (pelvis, bilateral thighs, shanks, and feet) from the standing calibration trial. The Pelvis Angle is typically defined as the orientation of the pelvis relative to the laboratory coordinate system, with 0° corresponding to the alignment of the pelvis coordinate system with the laboratory coordinate system. Knee joint angle will be zero when the thigh segment coordinate system and the shank segment coordinate system are aligned. The ankle joint angle is defined as 0° in the standing trial. The three-dimensional joint kinematics were calculated using an XYZ Cardan sequence of rotations, with X representing flexion-extension, Y representing abduction-adduction, and Z representing internal-external rotation (Wu et al. 2002). Shank inclination angle relative to the global vertical axis and GRF inclination angle relative to the global vertical axis are defined in the same way as Uno et al.(Uno et al. 2022).The joint moments were estimated using the inverse dynamics approach (Winter 2009). We normalized the joint resultant forces and moments by body weight. Each participant was represented by the ensemble average of five successful trials for each foot-strike condition.

### 2.5 Statistical analysis

Paired t-tests were used to evaluate the differences in kinematic and kinetic variables across various stop-jumping strategies. For the Statistical Parametric Mapping (SPM) analysis, paired t-tests were employed to compare two time-series data, examining the temporal variations in kinematic and kinetic measures across the stance phase. A custom MATLAB script was used to interpolate the data points into a time series curve consisting of 101 points, representing 0% to 100% of the landing phase. The statistical analysis was performed using the open-source SPM1d script for paired-sample t-tests to analyze the difference in kinematics and kinetics data during landing phase, with the significance level set at 0.001(Pataky, Robinson, and Vanrenterghem 2013).

## 3. Results

### 3.1 Joint angles in three planes (Sagittal, Frontal, Transverse)

Figure 4 shows the mean and standard deviation (SD) of right lower limb joint angles between initial foot contact and takeoff. In the sagittal plane, significant differences were found from 0.00% to 25.62% in the ankle joint angle (p < 0.001). In the frontal plane, significant differences were found from 39.59% to 45.59% in hip joint angle (p = 0.001); 16.17% to 29.42% in ankle angle (p < 0.001). In the transverse plane, significant differences were found from 0.00% to 31.12% in ankle joint angle (p < 0.001).

**Figure 4.**
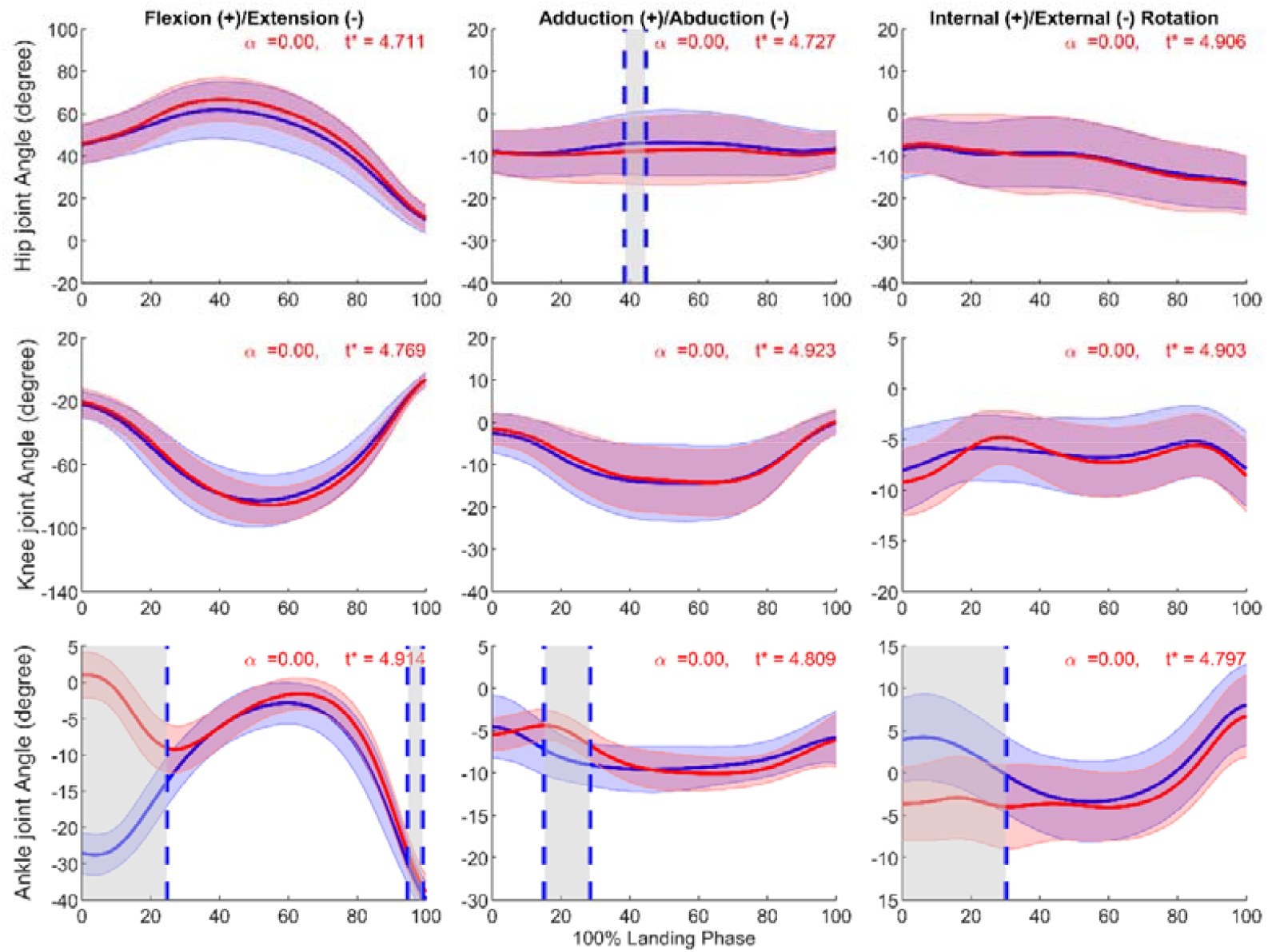
Mean±SD and SPM t-values of hip, knee, and ankle joint angles between forefoot landing (blue) and rearfoot landing (red) in three planes (Sagittal, Frontal, Transverse) from initial ground contact to takeoff (normalized to 100% landing phase) in 23 subjects. The significance level was set as p = 0.001. Statistical differences are highlighted in grey-shaded regions, indicating p < 0.001.

### 3.2 Joint moments in three planes (Sagittal, Frontal, Transverse)

Figure 5 shows the joint moment in the lower limb. In the sagittal plane, significant differences were found from 1.00% to 62.46% and 78.04% to 86.82% in ankle joint moment (p < 0.001). In the frontal plane, significant differences were found from 73.95% to 84.20% in ankle joint moment (p < 0.001). In the transverse plane, significant differences were found from 71.17% to 79.07% in hip joint moment (p < 0.001) and 0.00% to 3.56% in ankle joint moment (p < 0.001).

**Figure 5.**
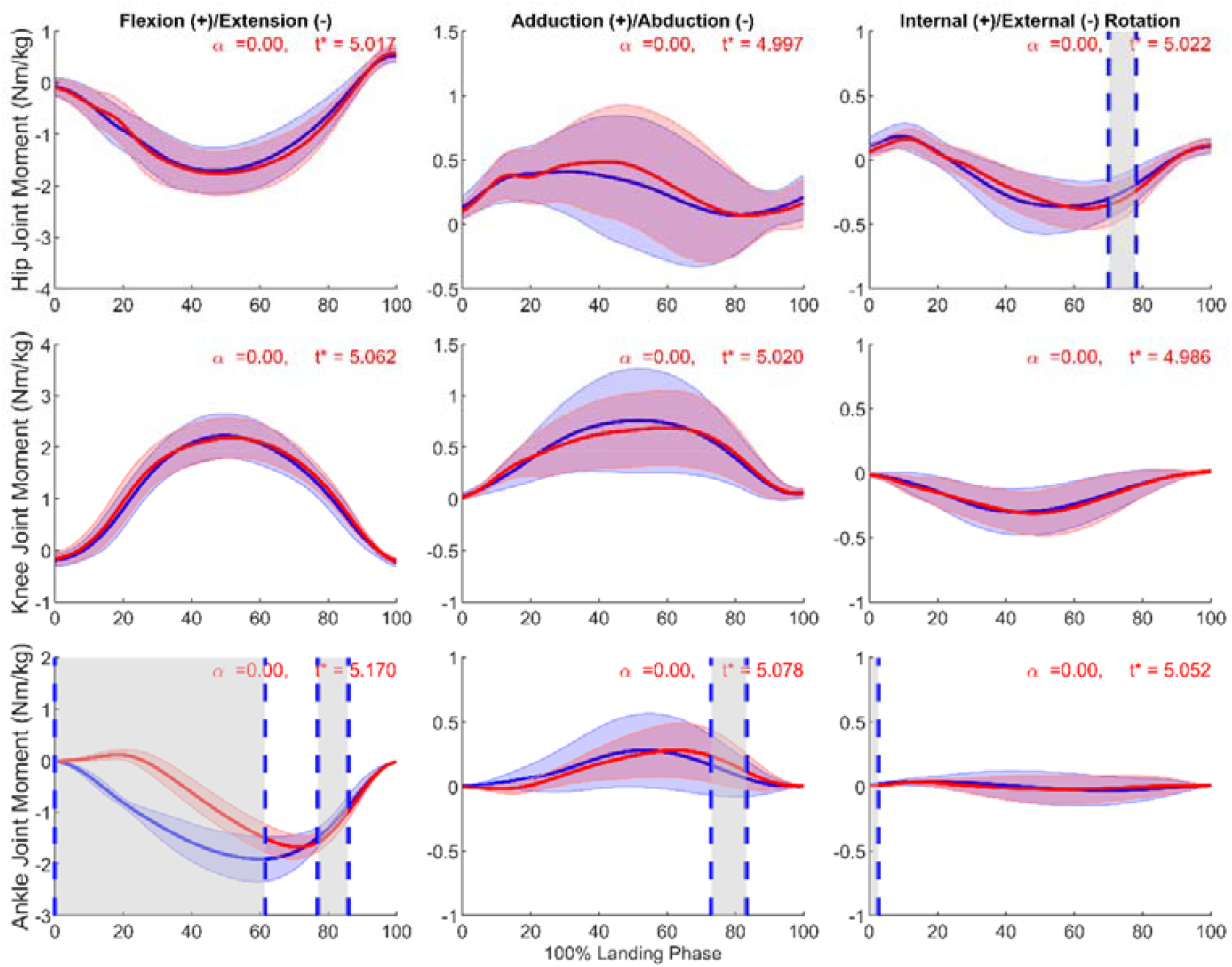
Mean±SD and SPM t-values of hip, knee, and ankle joint moments comparing forefoot landing (blue) and rearfoot landing (red) in three planes (Sagittal, Frontal, Transverse) from initial ground contact to takeoff (normalized to 100% landing phase) in 23 subjects. The significant level was set as p = 0.001. Statistical differences are highlighted in grey-shaded regions, indicating p < 0.001.

### 3.3 GRF, GRF inclination angle and shank inclination angle

Figure 6 shows the right lower limb GRF, GRF inclination angle, and shank segment inclination angle between initial foot contact and takeoff. In the anterior-posterior GRF, significant differences were found from 0.00% to 14.05% and 74.47% to 96.20% (p < 0.001). In the vertical GRF, significant differences were found from 75.46% to 83.02% in GRF (p < 0.001). In the sagittal plane, significant differences were found from 0.00% to 18.22% and 77.43% to 97.58% in GRF inclination angle (p < 0.001) and 18.5% to 46.56% in shank inclination angle (p < 0.001).

**Figure 6.**
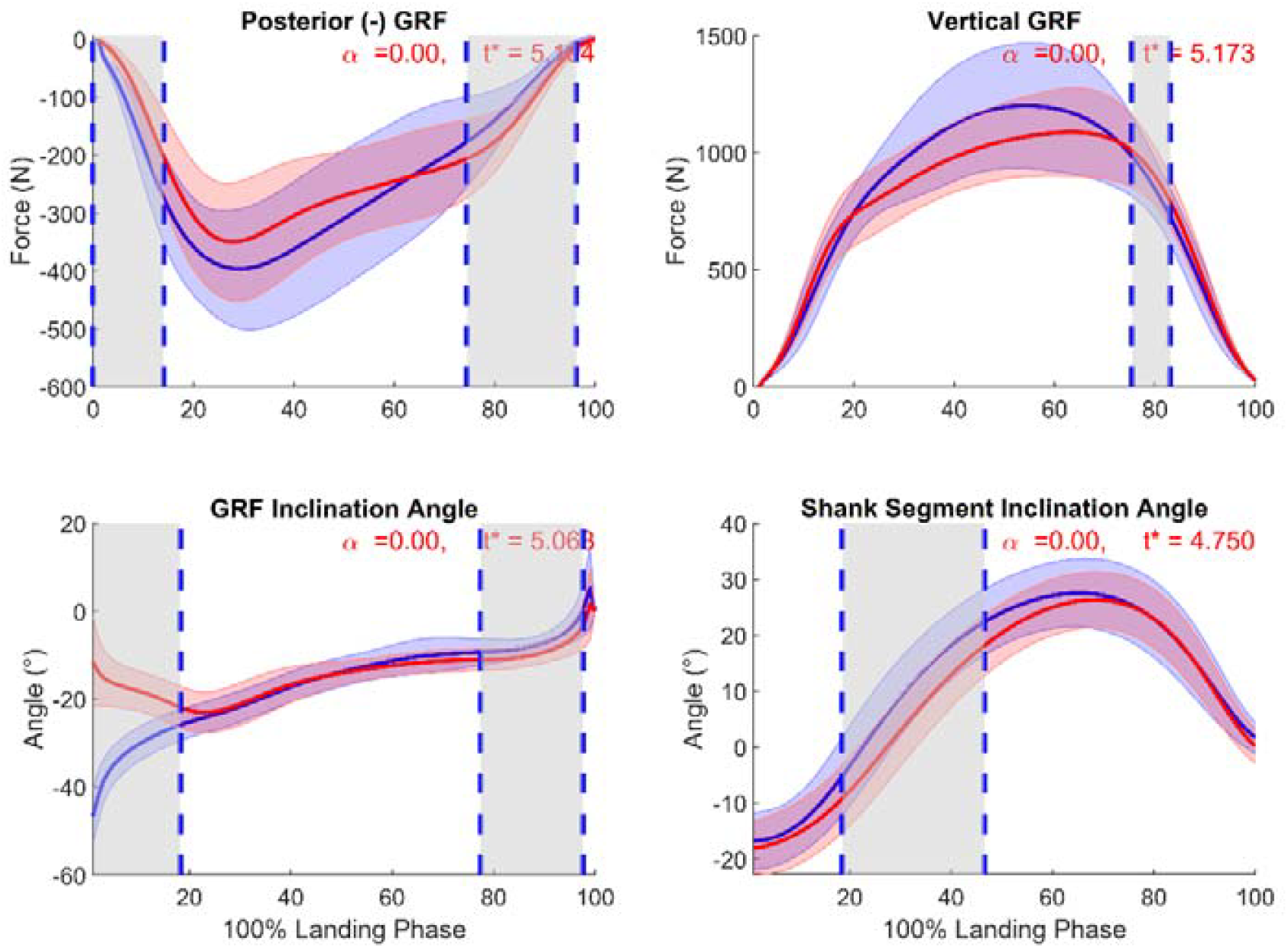
Mean±SD and SPM t-values of GRF in posterior and vertical direction, GRF inclination angle, and shank segment inclination angle comparing forefoot landing (blue) and rearfoot landing (red) from initial ground contact to takeoff (normalized to 100% landing phase) in 23 subjects. The significance level was set at p = 0.001. Statistical differences are highlighted in grey-shaded regions, indicating p < 0.001.

### 3.4 Ankle variables at initial contact

**Table 1.** Ankle variables at IC.

| Variables | Forefoot Landing | Rearfoot Landing | p-value | t |
| --- | --- | --- | --- | --- |
| Ankle angle at IC_X,<br>Degree | -26.02 (5.65) | 18.81 (5.61) | < 0.001 * | -18.86 |
| Ankle angle at IC_Y,<br>Degree | 3.45 (5.30) | 2.65 (3.06) | 0.56582 | 3.61 |
| Ankle angle at IC_Z,<br>Degree | 3.67 (4.59) | -3.32 (4.06) | < 0.001 * | 2.58 |
| Ankle joint moment at<br>IC_X, Nm/kg | -0.21 (0.06) | 0.03 (0.03) | < 0.001 * | -18.86 |
| Ankle joint moment at<br>IC_Y, Nm/kg | 0.02 (0.03) | -0.01(0.02) | 0.002 < 0.05 * | 3.61 |
| Ankle joint moment at<br>IC_Z, Nm/kg | 0.03 (0.02) | 0.02 (0.01) | 0.017 < 0.05 * | 2.58 |

Note: X, Y, Z represents sagittal, frontal, transverse plane separately. Nm: Newton\*meter. “\*” means significant difference. IC: initial contact.

In the sagittal plane (X), forefoot landing exhibited a significantly more plantarflexed ankle angle (−26.02° ± 5.65°) compared to rearfoot landing (18.81° ± 5.61°), with P < 0.001, and a normalized ankle joint moment (−0.21 ± 0.06 Nm/kg) that was significantly different from rearfoot landing (0.03 ± 0.03 Nm/kg), P < 0.001. In the transverse plane (Z), forefoot landing showed greater ankle angles (3.67° ± 4.59°) compared to rearfoot landing (−3.32° ± 4.06°), P = 0.001, and higher normalized ankle joint moments (0.03 ± 0.02 Nm/kg vs. 0.02 ± 0.01 Nm/kg), P = 0.017. No significant differences were observed in the frontal plane (Y) for ankle angles (3.45° ± 5.30° vs. 2.65° ± 3.06°, P = 0.56582), although forefoot landing showed slightly higher ankle joint moments (0.02 ± 0.03 Nm/kg vs. -0.01 ± 0.02 Nm/kg, P = 0.002).

### 3.5 Jump height, Stance time and Approach speed

**Table 2.** Mean (SD) of jump height, stance time and approach speed.

| Variables | Forefoot Landing | Rearfoot Landing | p-value |
| --- | --- | --- | --- |
| Jump height, cm | 49.51 (9.12) | 50.07 (9.06) | $0.34 > 0.05$ |
| Stance time, ms | 396.75 (100.27) | 433.48 (62.67) | $0.01 < 0.05^*$ |
| Approach speed, m/s | 2.14 (0.31) | 2.14 (0.34) | $0.85 > 0.05$ |

Note: cm: centimeter; ms: millisecond; m/s: miles per second. “\*” means significant difference.

Jump performance analysis revealed comparable jump heights between forefoot landing (49.51 ± 9.12 cm) and rearfoot landing (50.07 ± 9.06 cm, p = 0.34). Stance time was significantly shorter in forefoot landing (396.75 ± 100.27 ms) compared to rearfoot landing (433.48 ± 62.67 ms, p = 0.01). Approach speed remained consistent between techniques, with both forefoot and rearfoot landing showing identical speeds of 2.14 m/s (± 0.31 and ± 0.34 respectively, p = 0.85).

## 4. Discussion

The present study examined the effects of forefoot versus rearfoot landing strategies on lower limb biomechanics and performance during the stop-jump task. Our analysis revealed three key findings: (1) forefoot landing demonstrated increased ankle internal rotation during early landing, potentially elevating lateral ankle sprain risk; (2) forefoot landing showed biomechanical patterns that may reduce ACL injury risk; and (3) forefoot landing achieved comparable jump height with shorter stance time, suggesting potential performance benefits.

Our findings regarding ankle biomechanics support our primary hypothesis that forefoot landing would increase lateral ankle sprain risk. Previous investigators have demonstrated that ankle joint had more inversion and internal rotation angle comparing with normal status when lateral ankle sprain happened (Kristianslund, Bahr, and Krosshaug 2011; Fong et al. 2012; 2009). The increased inversion and internal rotation we observed during forefoot landing mirrors these injury-associated patterns and appears to be mechanistically linked to the lengthened moment arm of the ground reaction force around the subtalar joint (Shapiro et al. 1994; Barrett and Bilisko 1995). Specifically, the combination of plantar flexed, inversion and internal rotation landing positions observed in our study likely increases loading to the lateral ankle ligament.

Knee flexion angle at foot initial contact also has been regarded as a critical factor in NACL loading mechanics (Taylor et al. 2013; Yu and Garrett 2007). In our study, we found no significant differences in knee flexion angle between forefoot and rearfoot landing techniques during the initial contact phase, which is particularly relevant given that ACL injuries typically occur within 50ms after ground contact (B. Dai et al. 2015; Krosshaug et al. 2007, approximately 15% of landing phase in our study). This finding contrasts with Zhou et al., who reported significant differences in initial knee flexion angle during forefoot landing. Our results align more closely with Walsh et al. (2007), who found no significant differences in knee kinematics between controlled and soft landing techniques in male participants. The discrepancy between our findings and Zhou et al.’s study might be attributed to our emphasis on maximizing performance in the stop-jump task, as participants may have prioritized quick movement transitions over increasing knee flexion. This performance-oriented approach may have led participants in both landing conditions to maintain similar knee flexion angles that optimized their jump performance.

Our analysis of ground reaction forces and segment orientations revealed important differences between landing strategies. Consistent with Uno et al. (Uno et al. 2022)’s findings, forefoot landing demonstrated distinct characteristics: a larger posterior GRF component during the initial 14% of stance time and a more posteriorly inclined GRF angle in the first 18%. These biomechanical features may reduce ACL loading through two mechanisms: (1) decreased anterior drawer force from quadriceps activation, and (2) reduced proximal tibial anterior shear force. The tibial inclination angle provided additional insights into injury risk. While Boden et al. suggested that excessive compressive forces leading to anterior tibial translation are primary contributors to ACL injuries, our results showed that rearfoot landing produced greater posterior tibial inclination. This alignment is potentially problematic as it positions the femur to contact the relatively flat lateral region of the tibial plateau, potentially compromising joint stability and increasing injury risk.

Jump performance in sports requires both maximal height and quick execution, as minimizing stance time while maintaining jump height creates tactical advantages for subsequent movements. Our analysis revealed that forefoot landing achieved comparable jump heights with significantly shorter stance times compared to rearfoot landing. This faster ground contact time may be attributed to increased lower limb muscle activation both pre- and post-landing (Yoshida et al. 2016). While our findings appear to contrast with previous studies of forefoot-based soft landing techniques, this discrepancy likely stems from different methodological priorities. Specifically, soft landing studies typically emphasize impact reduction over performance optimization, leading to extended stance times (Boyi Dai et al. 2015). Our results suggest that forefoot landing can effectively balance performance demands, as evidenced by maintained jump height with reduced ground contact time, making it a potentially advantageous strategy for stop-jump tasks in competitive settings.

Several limitations should be considered when interpreting these results. Firstly, our exclusive focus on male subjects limits generalizability, particularly given that females have a higher incidence of noncontact ACL injuries than males (DeHaven and Lintner 1986; Lindenfeld et al. 1994; Arendt and Dick 1995; Ferretti et al. 1992). Secondly, fatigue could potentially influence the results of this study, as participants were required to complete multiple trials of each task at maximum effort. Nevertheless, we implemented several controls including randomized trial order and standardized rest periods to minimize fatigue effects. Third, our analysis was limited to the stop-jump task; future research should examine these landing strategies across other high-risk movements common in sports, such as cutting maneuvers and single-leg landings.

## 5. Conclusion

Our study demonstrates that forefoot landing during stop-jump tasks presents both advantages and risks. This strategy potentially reduces ACL injury risk and improves performance through decreased stance time without compromising jump height. However, the increased ankle internal rotation and inversion observed during forefoot landing may elevate the risk of lateral ankle sprains. These findings have important practical implications for athletes and coaches. While forefoot landing may be advantageous for performance and ACL injury prevention, its implementation in training programs should be accompanied by specific emphasis on developing ankle joint stability and control. Future research should examine whether targeted ankle stabilization training can help athletes maintain the performance benefits of forefoot landing while minimizing ankle sprain risk.

